# Hit or Miss: Understanding Emergence and Absence of Homo-oligomeric Contacts in Protein Language Models

**DOI:** 10.1101/2025.11.16.688745

**Authors:** Zhidian Zhang, Yo Akiyama, Yehlin Cho, Samarth Jajoo, Sergey Ovchinnikov

## Abstract

Many proteins function not as isolated molecules but as symmetrical assemblies of identical subunits, from ion channels that gate cellular signals to metabolic enzymes that catalyze life’s essential reactions. Here, we reveal that single-sequence protein language models (pLMs), trained solely on individual protein sequences, implicitly learn the interface contacts of homo-oligomeric assemblies. As models grow larger, their ability to predict inter-subunit contacts continues to improve, whereas the accuracy for single-chain predictions shows only minimal gains. The largest ESM2 model can accurately distinguish genuine biological interfaces from crystallographic artifacts. MSA Pairformer and ESM2-15B perform comparably when broad sets of homologs are used, but restricting alignments to closer evolutionary neighbors reveals a clear difference: MSA Pairformer reaches an interface contact recovery rate of 0.44 compared with 0.33 for ESM2-15B. We hypothesized that one contributing factor is how models implicitly cluster homologous proteins. Comparing evolutionary constraints extracted from pLMs, we find correlation between homologs decrease as model size increases, consistent with larger models partitioning families into finer-grained subclusters and better separating distinct oligomeric interfaces. When models fail to detect known interfaces, these discrepancies may reflect annotation errors, proteins that adopt multiple assembly conformations, or intrinsic model limitations. Overall, our findings show that statistical patterns learned by pLMs encode key aspects of homo-oligomeric assembly organization and provide a basis for understanding how such interfaces diversify across evolution, even though structure is conserved over large evolutionary distances, oligomeric assembly may not be.

## 1 Introduction

Many proteins do not function as single polypeptide chains. Instead, they assemble into homooligomers, complexes composed of identical subunits, to carry out tasks ranging from ion transport and signal transduction to metabolism and cytoskeletal organization (1). In these assemblies, the interaction between subunits governs cooperativity, allosteric regulation, and sensitivity to post-translational modifications or environmental factors such as pH (2). Disrupting these interfaces can cause disease (3), underscoring the importance of understanding not only the fold of an individual chain but also the interactions between identical chains.

Despite their ubiquity and functional importance, homo-oligomeric assemblies remain challenging to predict. Symmetry classifiers, such as Seq2Symm, can identify the overall arrangement of subunits, but they do not resolve residue-level contacts that define functional interfaces (4). Structure prediction methods such as AlphaFold2 (5) can model full quaternary structures, but the computational cost grows cubically with the number and length of subunits, limiting their use at the proteome scale. Moreover, even when high-resolution structures are available, distinguishing biologically relevant interfaces from crystallographic packing interfaces remains a persistent challenge for structure databases.

Recent advances in protein language models (pLMs) offer a promising route to understanding protein structure and function. Trained on millions of natural sequences without any protein structure information, these models nonetheless store evolutionary constraints encoding secondary and tertiary structure, amino acid substitution statistics useful for predicting effects of mutation due to conservation or coevolution, from sequences alone (6; 7). In principle, if evolution encodes the constraints required for subunit interactions, then pLMs should also contain information about these homo-oligomeric interactions. Direct-coupling analysis (DCA) showed that while most directly coupled residues are spatially close within a monomer, a subset of strongly coupled yet distant pairs often correspond to contacts across homo-oligomeric interfaces (8; 9), suggesting that quaternary constraints leave detectable traces in sequence statistics. Yet, it is unclear to what extent current pLMs capture such quaternary structure, how this ability scales with model size, and how it compares to methods that explicitly use multiple sequence alignments (MSA).

Here, we investigate the extent to which pLMs recover homo-oligomeric contacts and how this capability emerges with scale. We evaluate both single-sequence pLMs and an MSA-based method on their ability to predict inter-subunit contacts. We show that single-sequence models, trained only on individual protein chains, implicitly learn many of the residue–residue interactions at homooligomeric interfaces. As models grow larger, their inter-subunit contact prediction continues to improve even after single-chain contact accuracy has begun to plateau, and the best models can distinguish genuine biological interfaces from crystallographic artifacts with high accuracy. The compact MSA Pairformer (10) with far fewer parameters outperforms much larger single-sequence models when distant sequences are removed. Intriguingly, the cases where models “fail” to recover known interfaces often point to annotation issues and deeper biological insights (Fig. 1A), including misassigned biological assemblies and proteins that adopt multiple oligomeric states across evolution. Rather than introducing yet another specialized machine learning model, our work shows that existing models already encode rich information about homo-oligomeric assembly, and that carefully probing their successes and failures can both guide improvements to predictive methods and illuminate how homo-oligomeric interfaces diversify in nature.

**Figure 1.**
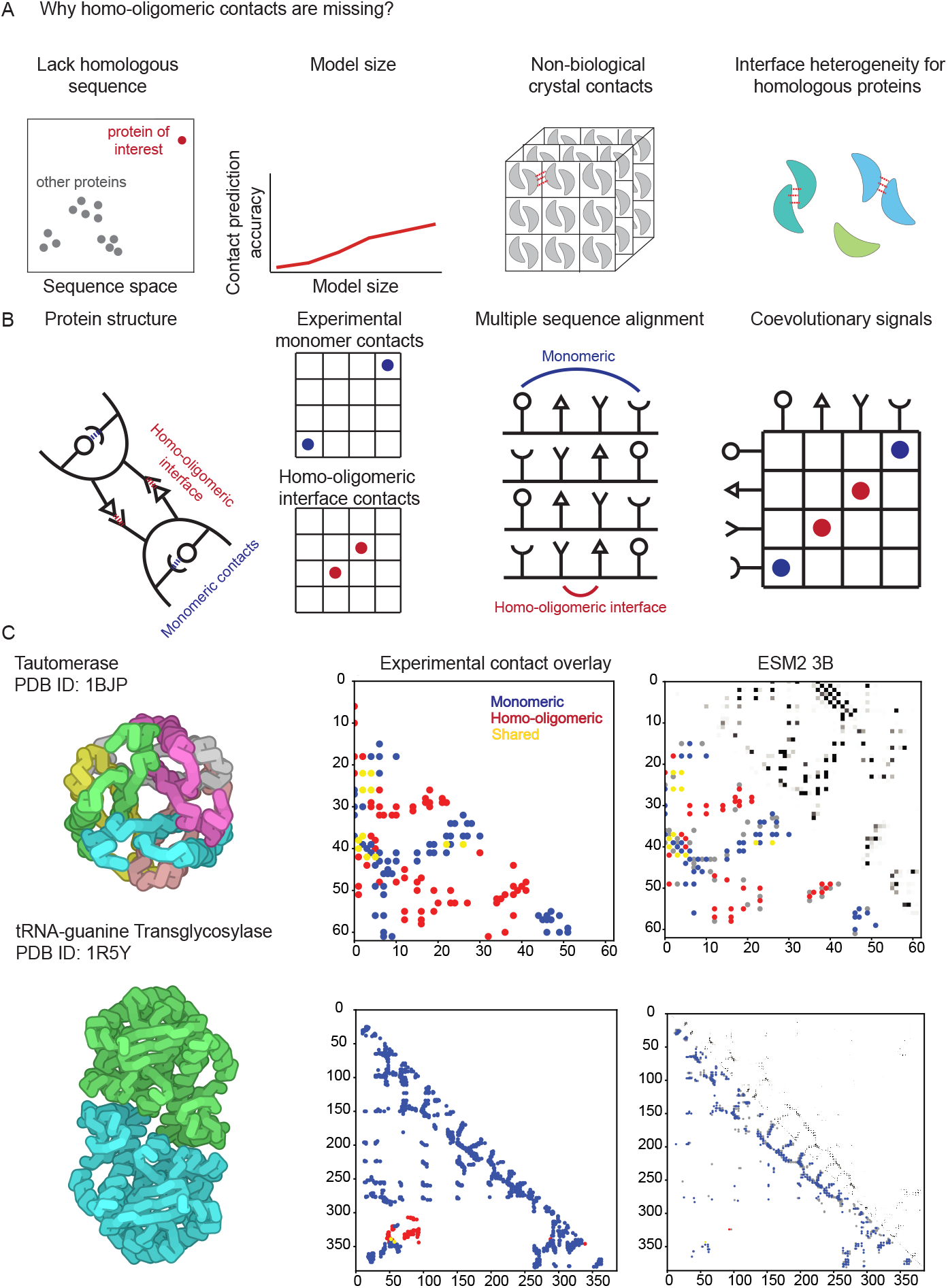
ESM2 recovers homo-oligomeric contacts from single-sequence input. (A) Conceptual overview of factors that can obscure homo-oligomeric interface signals, including limited homologous sequences, insufficient model capacity, crystallographic rather than biological contacts, and interface heterogeneity among homologs. (B) Homo-oligomeric proteins are composed of multiple copies of the same polypeptide. Coevolutionary signals captured by pLMs typically reflect a mixture of monomeric and homo-oligomeric interface contacts. (C) Tautomerase biological assembly (PDB ID: 1BJP), experimental contacts (sequence separation ≥ 6 residues), and ESM2-3B contact predictions showing matches to both monomeric and interface contacts. For tRNA-guanine transglycosylase (PDB ID: 1R5Y), ESM2-3B predictions lack homo-oligomeric interface signals.

## 2 Results

### 2.1 pLMs encode homo-oligomeric interface coevolutionary signals

Protein language models such as ESM2 (6) capture statistical dependencies imposed by evolutionary constraints. In these models, a simple logistic regression over attention maps is sufficient to predict residue–residue contacts within monomers (6; 11). Because evolutionary pressure acts at both tertiary and quaternary levels, we hypothesized that pLM attentions should also contain signals of homo-oligomeric interface contacts, in addition to monomeric contacts (Fig. 1B).

We illustrate this with tautomerase (Protein Data Bank PDB ID: 1BJP), a homo-hexameric enzyme (Fig. 1C). Providing only the monomer sequence to ESM2-3B yields a predicted contact map with prominent signals that cannot be explained by the monomer structure alone. After quantifying experimental monomeric and homo-oligomeric contacts (denoted as *X*) and taking the top *X* pLM contact predictions, we find that many of the top-scoring residue pairs that are distant within the monomer coincide with interface contacts in the biological assembly. This demonstrates that interface interactions can be recovered from single-sequence pLMs.

By contrast, for tRNA-guanine transglycosylase (PDB ID: 1R5Y), which also forms a homo-oligomer, the ESM2-3B predictions lack detectable homo-oligomeric signals (Fig. 1C). This discrepancy in homo-oligomeric contact recovery performance between the two experimentally resolved complexes motivated a systematic investigation of when pLMs capture homo-oligomeric contacts and when they fail to do so.

### 2.2 Homo-oligomeric signals are frequently missing

To quantify how often homo-oligomeric signals can be obtained from protein language models, we curated a benchmark of 4,160 nonredundant biological assemblies (30% sequence-identity clustering and Foldseek structure clustering; see Methods) (12; 13). For each assembly, we computed experimental monomeric and interface contact maps and evaluated ESM2 contact predictions by computing the precision at X, where X is the number of true contacts, which we term P@X (See methods).

Although tautomerase (PDB ID: 1BJP) displays a clear and strong interface signal, homo-oligomeric contact recovery is, on average, markedly lower than monomeric contact recovery across the dataset (ESM2-15B, Fig. 2A). Interface signals are therefore present but far from universal.

**Figure 2.**
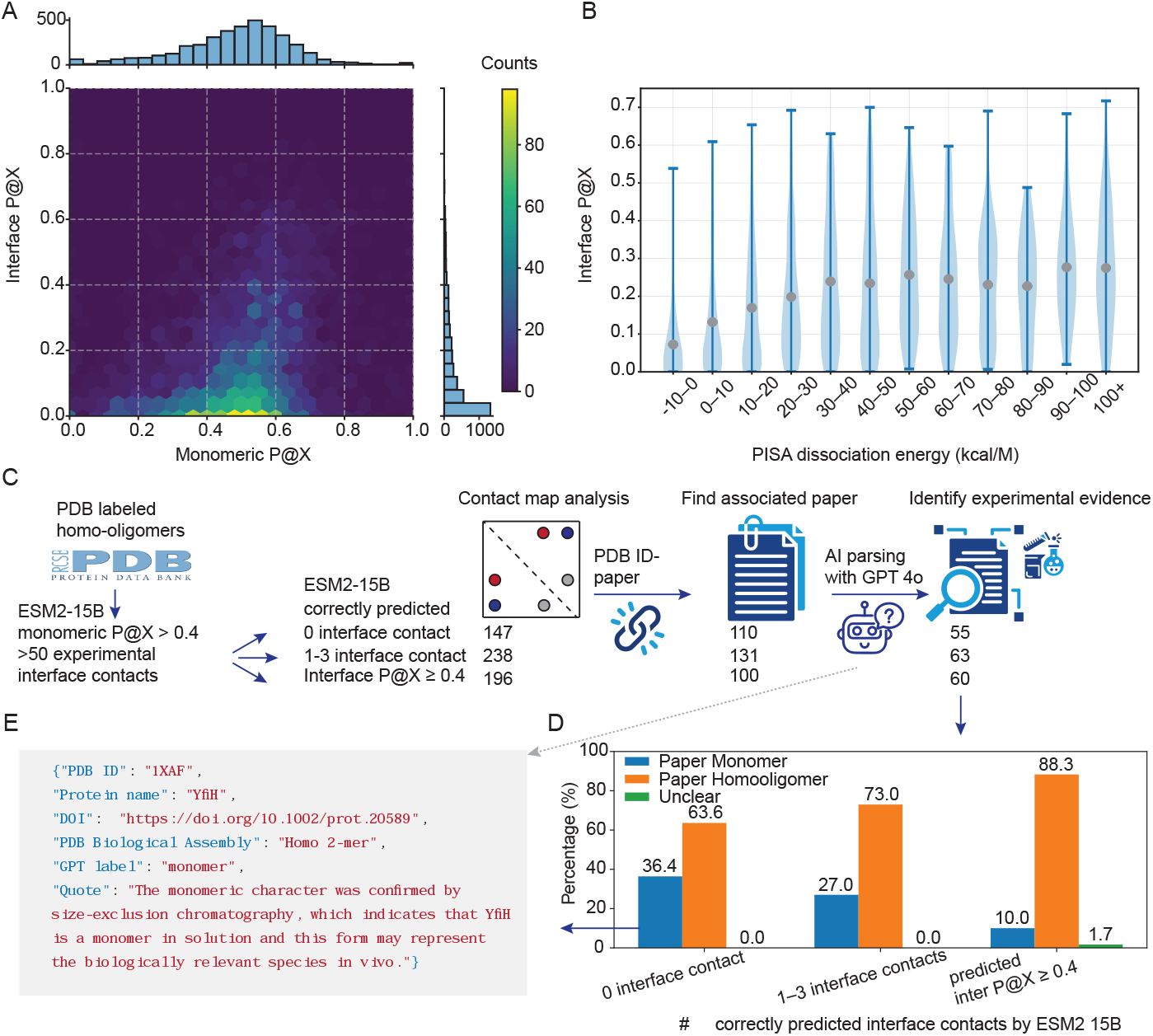
Why homo-oligomeric signals go missing. (A) Interface-versus monomeric-contact recovery for ESM2-15B across the nonredundant assembly dataset. (B) Interface contact recovery for biological assemblies with different PISA dissociation energies. (C) Workflow for extracting non-crystallographic experimental evidence of oligomeric state from primary publications. (D) Enrichment of solution-monomer reports in primary publications among PDB entries with poor interface contact signals compared to those with strong interface recovery. (E) Example of a mislabel: the primary paper reports a monomer in solution, whereas the PDB biological assembly is annotated as a homo-oligomer.

One hypothesis is that the interface strength correlates with the quality of the coevolutionary signal: obligatory, extensive interfaces might exhibit stronger covariation than weak or transient ones. Using the PDBePISA (Proteins, Interfaces, Structures and Assemblies, PISA) free energy of dissociation (the free energy change for separating the assembly into monomers) (14), we observe reduced interface contact recovery for assemblies with very low dissociation energy (Fig. 2B). However, substantial variance persists at all PISA dissociation energies, indicating that interface stability alone does not explain the observed heterogeneity in homo-oligomeric contact recovery

A second determinant is annotation quality. Although PDB biological assemblies are often treated as ground truth, we identified multiple cases in which solution experiments describe a monomer while the deposited biological assembly is oligomeric (and vice versa). To systematically assess this, we linked each PDB entry to its primary publication, expanded protein aliases using PDB and UniProt (15; 12), and applied a paper-parsing pipeline to extract non-crystallographic evidence of oligomeric state (e.g., size-exclusion chromatography with multi-angle light scattering, analytical ultracentrifugation, mass photometry; Fig. 2C and Methods).

Among structures with no detectable interface contact recovery by ESM2-15B (top-X evaluation; Methods), 20/55 primary publications explicitly reported the protein to be monomeric in solution. By contrast, among entries with high interface recovery (inter P@X ≥ 0.4), only 6/60 were described as monomeric (Fig. 2D). This enrichment suggests that missing interface signals often reflect genuine monomeric states or mislabeled biological assemblies rather than purely model failure. The YfiH structure (PDB ID: 1XAF) illustrates this: size-exclusion chromatography indicates a monomer, whereas the PDB biological assembly lists a homo-dimer; the absence of interface signal in ESM2-15B is consistent with the solution state (Fig. 2E).

### 2.3 Scaling pLM improves homo-oligomeric contact recovery

Because PDB biological assemblies can be noisy (misassigned), we further evaluated interface contact prediction on the manually curated EPPIC (Evolutionary Protein-Protein Interface Classifier) homo-oligomer benchmark (16) in which biological versus crystallographic interfaces are annotated (Methods).

Scaling ESM2 from 150M to 15B parameters improves both monomeric and interface contact recovery, but the relative gain in interface contact prediction exceeds that observed for monomeric contacts (Fig. 3A). Dehydroquinate dehydratase (PDB ID: 3O1N) exemplifies this pattern: interface contact signals are weak at 150M but emerge strongly at billion-parameter scales, whereas monomeric contact recovery is beginning to plateau at 150M (Fig. 3B). These observations suggest that larger models increasingly capture evolutionary constraints associated with quaternary structure.

**Figure 3.**
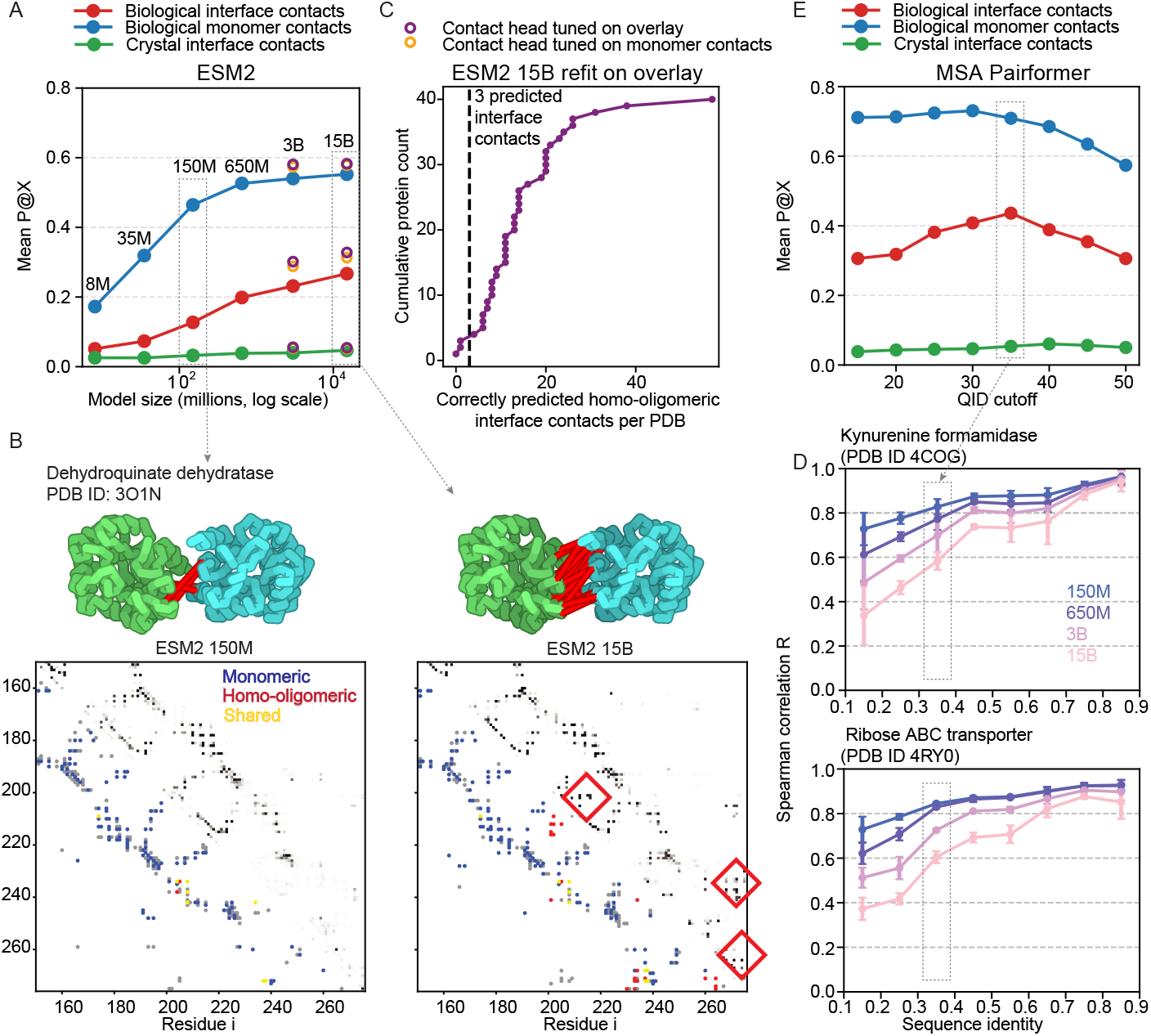
Scaling and removal of distant sequences improves homo-oligomeric interface prediction. (A) Homo-oligomeric interface and monomeric-contact recovery for ESM2 models of different sizes; models with the overlay-refit contact head are shown as circles. Crystal interfaces exhibit negligible pLM signals compared to biological interfaces. (B) Contact maps for dehydroquinate dehydratase (PDB ID: 3O1N) illustrate emergent interface signals at larger pLM scales. (C) Cumulative distribution of biological interfaces by the number of recovered interface residue pairs in ESM2-15B with the overlay-refit head; only 3/40 have fewer than three recovered interface pairs. (D) Spearman correlation between categorical Jacobians of aligned sequences and the query sequence shows that larger models yield more distinct categorical Jacobians for homologous sequences. (E) Homo-oligomeric interface and monomeric-contact recovery by MSA Pairformer versus MSA sequence-identity cutoff.

The original ESM2 contact head was trained on monomeric C*α* contact maps from 20 PDB structures, even though more than half of these proteins are homo-oligomers. To better align with what evolution encodes, we refit the contact head on an overlay of monomeric and homo-oligomeric contacts for proteins known to oligomerize (Methods), and evaluated contacts using ConFind (17) rather than a C*α*-distance cutoff, as ConFind contacts better reflect coevolutionary constraints (18). This refit improves both monomer and interface contact recovery (Fig. 3A).

A key application of homo-oligomeric signals from pLMs is discrimination between biological and crystallographic interfaces. On the EPPIC benchmark, which was curated so that biological and crystal interfaces are similar in size and therefore challenging to distinguish, pLM-derived contact signals are strong at biological interfaces but negligible at crystal packing contacts (Fig. 3A). Interestingly, for three entries in the crystallographic set (PDB IDs 1WLY, 1WOQ, and 2OW9), ESM2-15B predicts substantial interface contacts. Although the PDB author-assigned biological assemblies list these proteins as monomers, and the EPPIC dataset classifies these interfaces as crystallographic, EPPIC’s own interface classification server identifies assembly 1 as biological for all three, and PISA also assigns 1WLY and 2OW9 as homo-oligomers. These cases suggest that strong pLM interface signals may flag assemblies misclassified as monomers and support the existence of genuine biological homo-oligomeric interfaces.

Finally, although interface contact recovery is lower than monomeric recovery overall, only a small number of confident interface residue pairs are required to dock accurate monomeric structures into homo-oligomeric assemblies. Using ESM2-15B with the contact head refit on the overlay of experimental homo-oligomeric interface and monomeric contacts (Methods), we recover ≥ 3 interface contacts for 37/40 biological interfaces in the EPPIC benchmark (Fig. 3C), generally sufficient to constrain docking.

### 2.4 Homo-oligomeric interface heterogeneity among homologs attenuates interface signals

Why do some proteins with excellent monomeric contact recovery nonetheless lack interface contact signals in pLMs?

We hypothesized that one contributing factor is how models implicitly cluster homologous proteins. Larger models may build more refined clustering that separates subfamilies with distinct homooligomeric interfaces. To test this, we selected random protein families shorter than 300 residues with a wide range of sequence identities to the query and compared categorical Jacobians (19) for sequences in the MSA (Methods). Categorical Jacobian captures the evolutionary constraints on the sequence. Spearman correlations between the categorical Jacobians of the query and aligned sequences decreased as model size increased (Fig. 3D). For example, sequences with 30–40% identity to the query have Spearman *R* ≈ 0.8 for ESM2-150M and ESM2-650M, but only *R* ≈ 0.6 for ESM2-15B. This suggests that ESM2-15B distinguishes more sequence subclusters within the same family, consistent with larger models learning finer-grained clustering across homologs.

Single-sequence pLMs do not allow for explicit control of evolutionary context. To further investigate the extent to which homo-oligomeric contact prediction depends on subclustering within a protein family, we turned to MSA Pairformer, a compact 111-million-parameter model with state-of-the-art performance in monomeric contact prediction (10). With query-biased-attention turned off, this gives us full control over the evolutionary information used to make the prediction. Using MSAs constructed with HHblits and capped at a depth of 512 sequences (see Methods), we ran MSA Pairformer at different sequence-identity cutoffs to query (QID). ESM2-15B homo-oligomeric contact prediction is roughly comparable to using a minimally filtered MSA at 15% (Fig. 3E). Notably, we find that increasing the identity cutoff to 35%, and thus focusing on closer homologs increases the interface P@X from 0.33 to 0.44 (Fig. 3E). In typical MSA generation workflows, sequence depth is prioritized, and narrow sequence-identity ranges are rarely enforced. This strategy implicitly assumes that protein structures remain conserved across substantial evolutionary distances. Our results indicate that this assumption can fail for homo-oligomeric interfaces, which are often not conserved over such large evolutionary distances.

For proteins in the EPPIC dataset, 37/40 biological interfaces have at least three interface contacts recovered by ESM2-15B. The remaining three cases with the weakest ESM2-15B interface signals are 3EPW, 3CM3, and 2WXD. For 3CM3 (a dual-specificity protein phosphatase), domain swapping likely entangles interface and monomeric contacts, and another dual-specificity phosphatase (PDB ID: 2P4D) is reported to be monomeric in PDB, suggesting family-level heterogeneity. For 2WXD (a glucosidase), multiple related entries are reported as monomeric (e.g., 4PTW, 1UG6, 2E40, 2E9L), again pointing to mixed oligomeric states among homologs.

The inosine/uridine-preferring nucleoside hydrolase 3EPW highlights a different scenario: interface heterogeneity among proteins with similar monomeric folds. A related family member, 3MKM, forms a homo-tetramer with distinct interfaces despite a similar monomer architecture (Fig. 4A).

**Figure 4.**
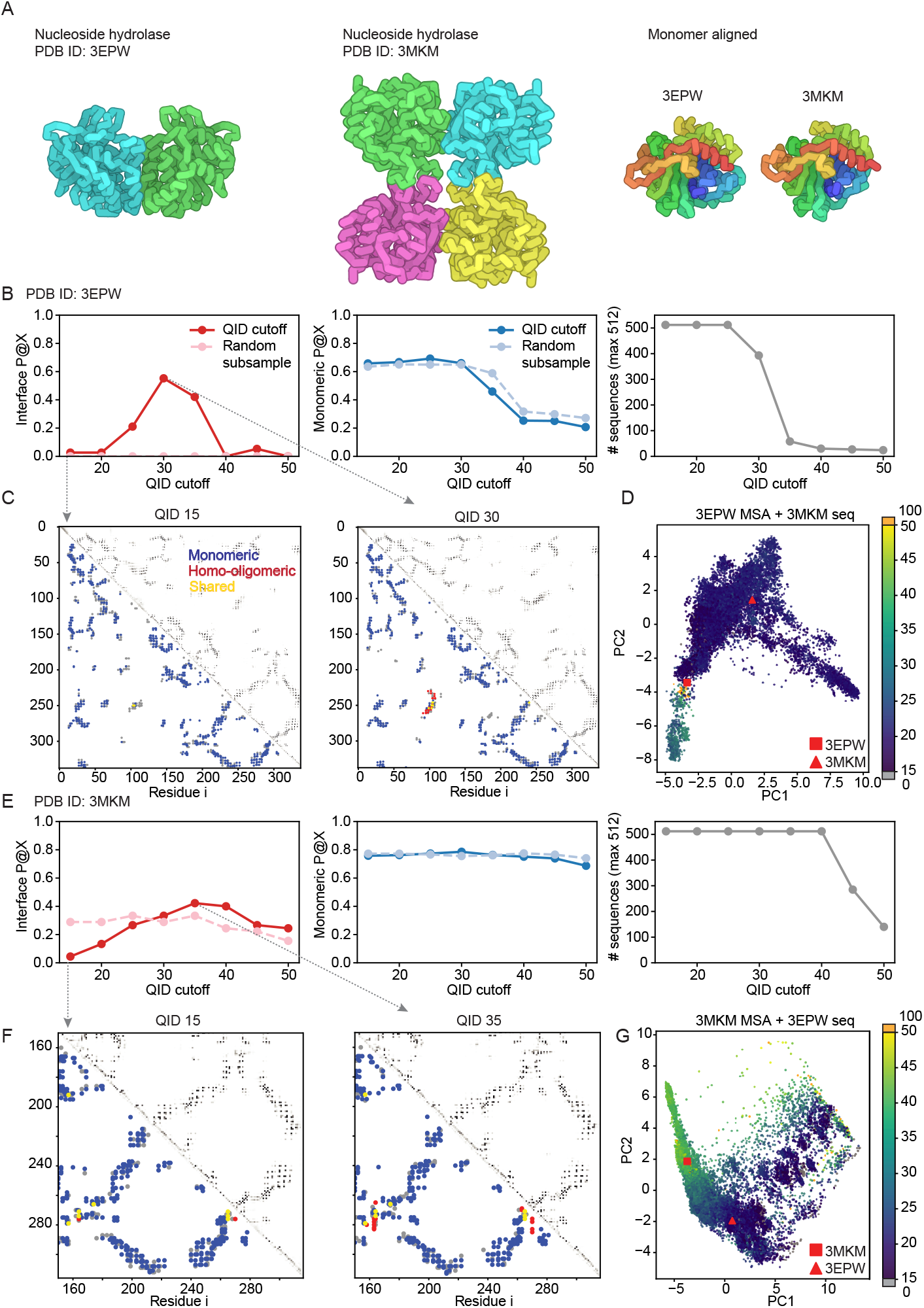
Homologous proteins with similar monomeric structures can have distinct homooligomeric interfaces. (A) Two members of the inosine/uridine-preferring nucleoside hydrolase (IU_nuc_hydro) family (3EPW, 3MKM) share similar monomeric folds but differ in homo-oligomeric contacts. (B,E) Interface and monomeric-contact recovery versus MSA sequence-identity cutoff (QID); matched-count random subsampling serves as a control. (C,F) MSA Pairformer contact maps at QID 15% and at the optimal QID. (D,G) PCA projections of MSA sequences colored by sequence identity to the query. The query and added sequence are indicated by square and triangle markers, respectively.

We systematically varied the MSA identity cutoff for MSA Pairformer (10). For both 3EPW and 3MKM, raising QID from 15% to 30–35% enriches the MSA for sequences closer to the 3MKM-like subfamily and increases interface contact recovery, whereas monomeric recovery remains essentially unchanged (Fig. 4B,C,E,F). We further combined the MSAs by adding the 3MKM sequence into the 3EPW MSA and vice versa, projected all sequences into a low-dimensional space using principal component analysis (PCA) (Fig. 4D,G). The two proteins occupy distinct sequence clusters, suggesting that the broader family contains subfamilies encoding different quaternary arrangements. When an MSA mixes such subfamilies, each with its own homo-oligomeric interface, the coevolutionary signal for any single interface can become diluted.

This behavior generalizes. Across families with ≥ 2 homo-oligomeric interface clusters (curated from ProtCID (20); Methods), scanning QID values above the default 15% frequently enhances interface contact recovery with minimal impact on monomeric contacts (Fig. 5). Matched-count subsampling controls show that these gains are not simply due to reducing MSA depth, but rather to selectively enriching interface-consistent sequences (Fig. 5).

**Figure 5.**
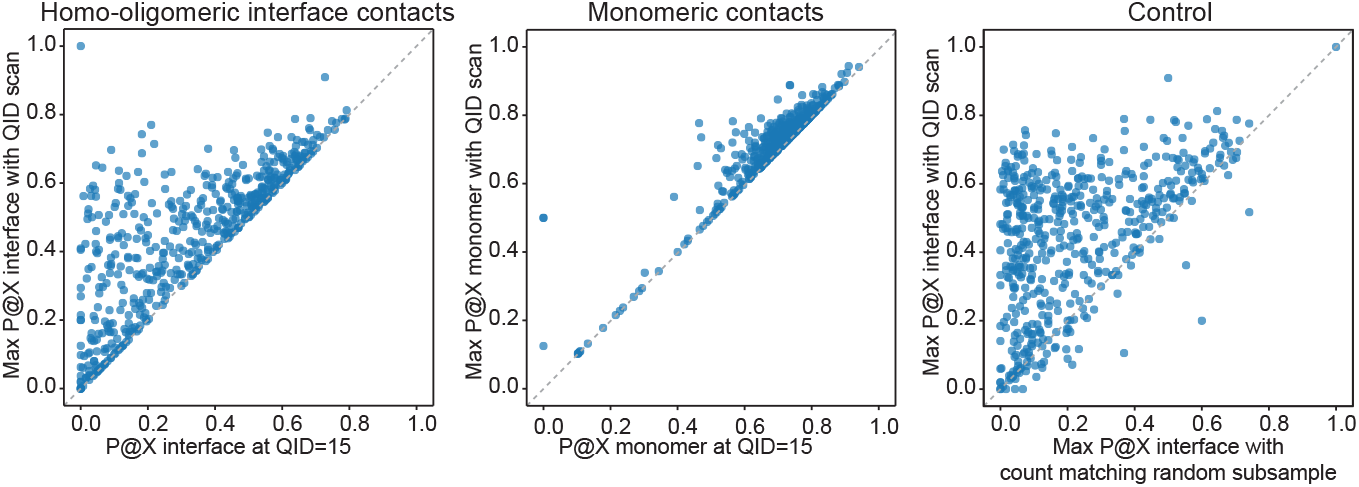
Removing distant sequences improves homo-oligomeric contact predictions. Left: Best interface contact recovery achieved by scanning different sequence identity to query cutoff (QID) values versus the default 15% QID. Middle: Best monomeric contact recovery achieved by scanning QID values versus the default 15% QID. Right: Best interface contact recovery achieved by QID scanning versus matched-count subsampling.

Together, these results suggest that interface heterogeneity among homologous sequences can dilute homo-oligomeric contact signals when MSAs mix subfamilies with divergent interfaces. Focusing on closer homologs (higher QID) restores the coevolutionary signature better. In the rare cases where random subsampling outperforms identity-based filtering, the similarity of homo-oligomeric interfaces may not correlate tightly with global sequence identity.

## 3 Discussion

Predicting quaternary structure remains a central challenge in structural biology. Methods such as AlphaFold2 have achieved impressive accuracy but are computationally demanding, especially for large assemblies and proteome-scale applications. Here, we show that pLMs, trained only on single sequences, already encode constraints that reflect homo-oligomeric organization. Interface contact signals extracted from model attentions reveal that the evolutionary information captured by pLMs extends beyond tertiary folds to homo-oligomeric quaternary assembly.

These interface signals arise because evolutionary selection acts on residue pairs that maintain both monomeric and homo-oligomeric interface interactions. Classic coevolution analyses have shown that residue pairs that co-vary yet are distant within a monomer can correspond to homo-oligomeric interfaces (8; 9). We demonstrate that this phenomenon persists in single-sequence models: attention-derived contact predictions contain signals corresponding to known interface residue pairs, even without multi-chain input. Quaternary constraints are thus embedded within the same sequence statistics that govern monomeric structure.

Across a nonredundant benchmark, interface contact recovery is systematically lower and more heterogeneous than monomeric contact recovery. This variability reflects both technical and biological factors. First, many PDB biological assemblies annotated as oligomeric correspond to monomers in solution, meaning that apparent “failures” to predict interfaces may in fact reproduce the true monomeric state. Second, the intensity of the evolutionary signal depends on interface stability: transient or condition-dependent assemblies leave weaker, less consistent covariation patterns. Third, interface heterogeneity among homologous sequences dilutes subfamily-specific couplings. When MSAs mix proteins that share a monomeric fold but differ in oligomeric state or interface, the signal for any one interface is averaged away. Focusing MSAs on interface-consistent sequences, by increasing the sequence-identity cutoff, systematically enhances interface contact signal without degrading monomeric contact recovery. This aligns with previous results from DCA: for response-regulator homo-oligomers, interface contact prediction accuracy improves when family subtypes with distinct binding modes are analyzed separately (9; 21).

Model scaling markedly strengthens homo-oligomeric interface signals. As pLMs increase from hundreds of millions to tens of billions of parameters, homo-oligomeric interface contact recovery improves, often more than monomeric contact prediction. Analysis of categorical Jacobians suggests a mechanistic explanation: larger models form more finely partitioned sequence clusters, effectively grouping homologs with similar oligomeric behaviors. Such refined clustering likely sharpens the model’s internal separation between distinct homo-oligomeric interface-specific couplings. In turn, this enables large pLMs to better discriminate among homologous proteins that share a common monomeric fold but assemble through different interfaces.

The interface signals extracted from pLMs also provide a powerful discriminator between biological and crystallographic contacts. On the EPPIC benchmark (16), pLM-derived contacts are abundant at biological interfaces yet largely absent at crystal-packing contacts, even when the two classes are matched in size. In several cases where ESM2 predicts strong interface signals for “crystallographic” contacts, independent evidence from EPPIC and PISA supports these interfaces as biological. These results suggest that coevolutionary information from pLMs can complement physics-based metrics such as buried surface area or thermodynamic stability, providing an orthogonal and scalable criterion for identifying biologically relevant assemblies and flagging misannotated structures.

Several limitations remain. Evolutionary data average over cellular and environmental contexts (22), so interfaces that form only under specific conditions may not leave strong or consistent sequence signatures. Our current MSA processing also relies on heuristic sequence-identity thresholds, whereas more systematic subfamily clustering could further sharpen interface signals.

Despite these caveats, the practical implications are immediate. A single forward pass through a pLM can often identify enough interface residue pairs to dock monomeric structures into accurate homo-oligomeric assemblies. This lightweight strategy provides a scalable alternative to multi-chain inference by models such as AlphaFold2 (5) and enables rapid screening of candidate assemblies in metagenomic datasets, structural genomics efforts, and annotation of existing PDB entries.

Our findings also align with an emerging view of quaternary structure evolution as a mix between functional selection, entrenchment, and neutral drift. Some interfaces become entrenched through hydrophobic ratchets (23), remaining fixed even when no longer strictly required for function, while others transition repeatedly between oligomeric states without major impact on activity (24). The variable intensity of interface contact signals observed across protein families likely reflects this evolutionary heterogeneity: strongly conserved, functionally entrenched assemblies produce robust pLM interface signals, whereas labile or context-dependent oligomers might leave only faint traces in sequence.

In summary, large pLMs inherently encode homo-oligomeric interactions: the statistical patterns they learn from sequences alone capture not only how individual chains fold, but also how they assemble into oligomers. By extracting interface contacts from large single-sequence pLMs or compact MSA-based pLMs, we can recover biologically relevant homo-oligomeric interfaces, discriminate them from crystal packing interfaces, and even identify assemblies that are likely misannotated. The most intriguing cases are those that appear as “failures” of prediction, where weak or conflicting interface signals instead highlight true monomeric states or families that toggle between distinct oligomeric configurations.

## 4 Methods

### 4.1 Data Collection and Filtering

We began with 326,530 macromolecular structures from the Protein Data Bank (PDB) (12) as of 10/01/2024. Homo-oligomeric stoichiometry information was retrieved for 225,681 entries on 10/02/2024, yielding 76,801 candidate homo-oligomers.

To ensure data quality and simplify downstream processing, we applied the following filters:

- **Copy number**: Complexes with ≥10 identical chains were excluded.
- **Resolution**: Structures with resolution *>*5 Å were removed.
- **Chain length consistency**: All chains within a complex were required to differ in length by no more than 10%, and each chain had to contain 20–500 resolved residues.
- **Structural completeness**: Assemblies with gaps of *>*50 consecutive unresolved residues in any chain were excluded.

After filtering, 58,920 high-quality homo-oligomeric assemblies remained. To reduce redundancy, we clustered sequences at 30% identity using PDB UniRef clusters (9,762 clusters), and further clustered structures using Foldseek, yielding 4,309 structural clusters. From each Foldseek cluster, we selected a representative structure. We downloaded the bio-assembly1 for each structure.

Assemblies with ambiguous residue numbering in the Macromolecular Crystallographic Information File (mmCIF) files, ConFind (17) processing failures, or chain lengths exceeding 500 residues were excluded, leaving 4,160 assemblies.

### 4.2 PISA and literature analysis data preparation

We began with 4,160 homo-oligomeric assemblies. To focus on interfaces large enough for meaningful analysis and on proteins for which the language model accurately captures monomeric structure, we retained assemblies with more than 50 predicted interface contacts and monomeric P@X *>* 0.4. This filtering yielded a curated set of 1,583 assemblies. Precomputed interface parameters were retrieved from PDBePISA (Proteins, Interfaces, Structures and Assemblies) for 1,291 of the 1,583 assemblies.

### 4.3 Paper parsing with GPT application programming interface (GPT API)

We programmatically extracted oligomerization evidence from publications for selected PDB structures. First, we assembled DOIs for PDB IDs that met an a priori filter based on ESM2 predictions (0 predicted interface contacts, or 1–3 predicted interface contacts, together with interface contact recovery ≥ 0.4). Full-text PDFs were parsed with PyMuPDF to extract relevant sentences. To avoid cross-assignment in multi-structure papers, we detected and excluded sentences that mentioned non-target PDB codes, and expanded protein aliases using PDB metadata (12) and UniProt (15) to increase recall.

Sentences were scored with a keyword/section heuristic that prioritizes quantitative, solution-state methods including size-exclusion chromatography (SEC), SEC coupled with multi-angle light scattering (SEC-MALS), analytical ultracentrifugation (AUC), native mass spectrometry (native MS), blue-native polyacrylamide gel electrophoresis (BN-PAGE), chemical cross-linking, small-angle X-ray scattering (SAXS), and mass photometry. The top-ranked context was passed to a large language model (OpenAI GPT-4o) constrained by a strict JSON schema to return: (i) yes/no for homo-oligomer status, (ii) whether evidence was inferred only from crystallography, and (iii) quotations. A protein is classified as a homo-oligomer only if at least one extracted item explicitly supported homo-oligomerization by a biophysical method; reports based solely on crystallography were labeled as “inferred only”. All per-PDB outputs were written to JSON for downstream verification and analysis. We started with 147, 238, and 196 pdb entries in each category (0 predicted interface contacts, 1–3 predicted interface contacts, interface contact P@X ≥ 0.4 by ESM2-15B). We successfully retrieved 110, 131, and 100 papers, of which 55, 63, and 60 contained non-crystallographic biophysical evidence of oligomerization.

### 4.4 Evaluation on the EPPIC crystal and biological interface dataset

We downloaded the Duarte-Capitani crystal interfaces dataset (DCxtal) and biological interfaces dataset (DCbio) from www.eppic-web.org (16) on Aug 20th 2025. DCxtal comprises crystal contacts with interface areas >1000Å^2^, while DCbio contains biologically interfaces with interface areas <2000Å^2^. Entries with multiple reported interfaces or missing interface contacts according to our ConFind (17) cutoff were excluded, yielding 40 biological assemblies and 75 crystal assemblies.

### 4.5 Precision at X (P@X) Calculation

To evaluate contact prediction performance, we adapted the Precision@L metric to assess interface and monomeric contact recovery. Contacts were defined using ConFind (17), where residue pairs with contact degree *>* 0.01 and sequence separation ≥ 6 were considered. We chose to use ConFind contact degree for evaluation rather than C*α* distance because it aligns better with coevolutionary information.

Let *X* denote the total number of ground-truth contacts, either interface or monomeric contacts. From the predicted contact map, we extract the top *X* highest-scoring residue pairs (with sequence separation ≥ 6). Precision at *X* was computed as:

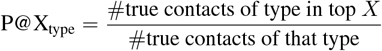

where *type* refers to one of the following:

- **interface-exclusive**: interface and not monomeric contacts,
- **monomeric-exclusive**: monomeric and not interface contacts

This formulation allows us to evaluate model performance for each contact type while accounting for label overlaps.

### 4.6 Refit contact head

Experimental contact maps were computed using ConFind (17), with a contact-degree cutoff of 0.01. We refitted the ESM2 (3B and 15B) contact heads on either (i) experimental monomeric contacts only, or (ii) an overlay of monomeric and interface contacts. For the 4,160 assemblies, we excluded sequences belonging to the same PDB 30% sequence-identity cluster as the EPPIC (16) dataset to prevent data leakage, and fit on the remaining assemblies.

### 4.7 Query identity cutoff screening

We used ProtCID (20) to identify protein families with multiple homo-oligomeric interface clusters. Clusters containing at least five PDB and PISA entries were retained, yielding 411 protein families. For each family, we sampled one PDB structure from each of the two largest interface clusters. Families with processing errors were excluded, resulting in 736 structures.

Multiple sequence alignments (MSAs) were constructed using HHblits (25) (three iterations, *E*-value cutoff 1 *×* 10^−5^, minimum coverage 0.75), followed by HHfilter (26) with sequence-identity cutoff 90% and query-identity (QID) range 15–50%.

Contact maps were predicted using MSA Pairformer (without query bias) at each QID cutoff, using a maximum sequence depth of 512. We analyzed 537 assemblies from the benchmark set (17); four entries were excluded due to sequence–structure mismatches caused by His-tags, and the remaining excluded cases were monomeric in PDB biological assembly 1. As a control, we repeated all analyses using random subsampling to match the sequence counts at each QID cutoff.

### 4.8 Spearman correlation of categorical Jacobians between MSA sequences

The categorical Jacobian (19) is a mean-centered and symmetrized tensor with size [*L, A, L, A*]. *L* is the length of the protein and *A* is 20 for the 20 different amino acid types. The categorical Jacobian indicates the coevolutionary statistics between all residue pairs learned by the protein language models.

Given the categorical Jacobian **J**(*X*) of a protein language model for a sequence *X* (as defined in (19)), and the corresponding language-model contact map **C**(*X*) derived from **J**(*X*), we quantified how similar the categorical Jacobians are for two homologous sequences drawn from the same MSA.

For each protein family, we took the MSA query sequence as the *reference* and selected additional sequences from the same MSA as *homologs*. For each reference–homolog pair, we first restricted the analysis to the aligned residues. After this filtering, the effective sequence length was *L*.

For each reference–homolog pair we obtained:

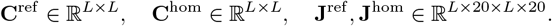

We then considered all residue pairs in the upper triangle of the *L × L* contact map, to avoid double-counting symmetric pairs.

To focus on residue pairs that the language model predicts to be strongly coupled, we ranked all residue pairs (*i, j*) by the reference contact score 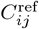 in descending order and retained only the top *L* pairs. For these *L* pairs we collected the corresponding 20 *×* 20 categorical Jacobian blocks from the reference and homolog sequences, yielding tensors

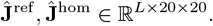

Because many Jacobian entries are close to zero and not informative, we further restricted the comparison to Jacobian entries with relatively large magnitude in the reference. Specifically, we computed a root-mean-square magnitude over all entries of 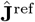, and selected entries with absolute value larger than this threshold.

Finally, we flattened filtered entries of the reference and homolog Jacobians into two vectors of equal length and computed the Spearman rank correlation between them, using the implementation in SciPy (27). This quantifies how similar the model’s categorical Jacobians are between the MSA query sequence and a homolog in the MSA. (19).

## 5 Code availability

All code used for data processing, model evaluation, and analysis will be released at: https://github.com/zzhangzzhang/homo_oligomer_contacts

## 6 Acknowledgment

We thank Roland Dunbrack and Qifang Xu for assistance in curating the ProtCID-derived dataset, and Roshan Rao, Gaetano T Montelione, and György Abrusán for valuable discussions. Z.Z. acknowledges the funding from the Swiss National Science Foundation Postdoc.Mobility fellowship [P500PB_225547] and MIT-Novo Nordisk Artificial Intelligence Postdoctoral fellowship. S.O. acknowledges funding from NSF grant MCB2032259, Coca-Cola and Amgen.

